# A nematode retrotransposon in the common shrew: horizontal transfer between parasite and host

**DOI:** 10.1101/424424

**Authors:** Sonja M. Dunemann, James D. Wasmuth

## Abstract

Reports of horizontal transposon and gene transfers involving metazoan species has increased with the sequencing of their genomes. Horizontal transfer could be facilitated by the intimate relationship between a parasite and its hosts. To date, two studies have identified horizontal transfer of RTEs, a class of retrotransposable elements, involving parasites: ticks might act as vector for BovB between ruminants and squamates, and AviRTE was transferred between birds and parasitic nematodes. We wanted to know if parasitic nematodes are involved in other cases of horizontal transfer of RTEs. We searched 33 mSammalian RTEs in 81 nematode assemblies, and 10 nematode RTEs in 98 mammalian assemblies. We identified RTE1õ Sar from *Sorex araneus*, the common shrew, in parasitic nematodes and show that it originates from nematodes. To exclude contamination of the *S. araneus* assembly, we developed an approach that uses long reads and paired-end reads. With phylogenetic analysis and copy age estimation, we show that RTE1_Sar was horizontally transferred from nematodes to *S. araneus*. We confirm horizontal transfer of RTEs in host-parasite interactions, and we present a new method to distinguish between contamination and horizontal transfer.

## Introduction

Transposable elements, or transposons, are vertically transmitted in eukaryotes. In some cases however, a transposon might be horizontally transferred to a different species. BovB (Bovine-B), a retrotransposable element present in ruminants and squamates, has been extensively studied as first case of a horizontal transfer of a retrotransposable element between eukaryotes [1, 2, 3]. Recently, two other cases of horizontal retrotransposon transfer have been reported: AviRTE has been detected in birds and human parasitic nematodes [4], and LINE-1 has possibly been transferred between marine eukaryotes [3].

It has been hypothesized that transposons survive longer if they acquire a new host through horizontal transfer by escaping host silencing mechanisms [5]. Although the mechanisms of horizontal transposon transfers (HTTs) are still unclear, they could be facilitated by parasites and pathogens [5, 6, 7]. Parasites have long lasting, physical contact to their hosts, which increases the chances of HTT either directly or through a secondary pathogen. The ectoparasitic ticks and bed bugs and their hosts have horizontally transferred BovB [2] and SPIN [6], respectively. Examples from endoparasites include AviRTE transferred between filarial nematodes and birds [4], and the transposon hsmar1 shared between hookworms humans [8].

After being horizontally transferred, transposons can have significant impact on the recipient organisms. Transposons proliferate in the host genome and increase the genome size. Transposons and their intra-genomic movement lead to changes in a species’ genotype and phenotype (reviewed by [9, 10]). Inter-species transfer of transposons have been shown to support the creation of new genes [11].

It is important to distinguish horizontal transfer from contamination of genetic material added during the sequencing process. This is not a simple task but necessary if we are to understand if and how frequent horizontal transfer occurs and what evolutionary forces shape these events. Contamination has been previously erroneously identified as horizontal transfer in the literature [12]. Given the close physical and molecular association between a parasite and its host, determining horizontal transfer is greater challenge.

Given the previously documented horizontal transfers of BovB and AviRTE, are RTEs more prone to horizontal transfer between parasites and hosts than between free-living species? To answer this question, it is necessary to comprehensively survey the rates of horizontal transmission for all types of genome elements across the tree of life. To this end, we investigated mammalian and nematode RTEs to identify potential transfers between parasitic nematodes and their hosts. We identified RTE1_Sar in nematodes and *S. araneus*, and BovB from cattle and sheep in nematodes. BovB has been analyzed previously in multiple studies, but has never been reported in nematodes, which lead to the question of whether the presence of RTE1_Sar and BovB resulted from misassemblies due to contamination from/to endoparasites. To exclude contamination, we searched for contamination of other genomic elements in *S. araneus* from the same nematode species that harbour RTE1_Sar, but found none. For *H. contortus*, long reads were available which we exploited to distinguish between HTT and contamination, and found that BovB is a contaminant. We then aimed to develop a similar read-based approach with short, paired-end reads for RTE1_Sar in *S. araneus*. We confirmed horizontal transfer of AviRTE in birds and nematodes with the long reads approach, and used AviRTE as positive control and BovB as negative control for the short reads approach. The short read-based approach takes advantage of paired-end reads, where only one of the read mates codes for a transposon. This approach was successful, and confirmed that RTE1_Sar in *S. araneus* is endogenous. We then tested for horizontal transfer of RTE1_Sar between *S. araneus* and nematodes by estimating a phylogeny and relative RTE1_Sar copy ages. We found that RTE1_Sar was horizontally transferred between an unsampled parasitic nematode and *S. araneus* after the split of the lineages leading to Soricinae and Erinaceus ca. 64 million years ago (mya) [13]. Our results emphasize the importance of checking for contamination as a spurious indicator of horizontal transfer, but also reveal novel cases of HTT.

## Results

### Horizontal transfer of RTEs between parasitic nematodes and their hosts is not common

To detect potential cases of horizontal transfer of RTEs between parasitic nematodes and their hosts, we performed reciprocal sequence similarity searches. We used BLAST to compare 33 mammalian RTEs (RepBase2014 [14]) to 81 nematode genomes (WormBase ParaSite 6 [15]), and in addition 10 nematode RTEs to 98 mammalian genomes (RefSeq [16], List of RTEs and genomes in Supplemental Files 1 and 2). To reduce false positives, at least 10 RTE copies were necessary per species.

We detected two RTEs, RTE1_Sar from the common shrew (*S. araneus*) and BovB from cattle and sheep (*Bos taurus*, *Ovis aries*), in nine and two nematode species, respectively (Table 1). All of the identified nematode species are parasites of mammals. RTE1_Sar-containing nematodes *Nippostrongylus brasiliensis* and *Angiostrongyloides cantonensis* have been found in the asian house shrew *Suncus murinus* [17], which, with *Sorex*, is a member of the family Soricidae.

**Table 1:** Mammalian RTE elements identified in nematode genome assemblies

| RTE | Hits | Nematode | Accession |
| --- | --- | --- | --- |
| BovB | 416 | *Haemonchus contortus* | PRJEB506 |
| BovB | 77 | *Onchocerca ochengi* | PRJEB1204 |
| BovB_Oa | 194 | *Haemonchus contortus* | PRJEB506 |
| BovB_Oa | 385 | *Onchocerca ochengi* | PRJEB1204 |
| RTE1_Sar | 155 | *Angiostrongylus costaricensis* | PRJEB494 |
| RTE1_Sar | 317 | *Angiostrongylus cantonensis* | PRJEB493 |
| RTE1_Sar | 40 | *Ancylostoma ceylanicum* | PRJNA231479 |
| RTE1_Sar | 26 | *Ancylostoma ceylanicum* | PRJNA72583 |
| RTE1_Sar | 48 | *Haemonchus contortus* | PRJEB506 |
| RTE1_Sar | 12 | *Haemonchus placei* | PRJEB509 |
| RTE1_Sar | 68 | *Heligmosomoides polygyrus* | PRJEB1203 |
| RTE1_Sar | 27 | *Necator americanus* | PRJNA72135 |
| RTE1_Sar | 56 | *Nippostrongylus brasiliensis* | PRJEB511 |
| RTE1_Sar | 43 | *Teladorsagia circumcincta* | PRJNA72569 |

### Contamination in the *S. araneus* genome assembly does not confound RTE1_Sar HTT

Contamination of genome assemblies is common and must be considered when analyzing for HTT. To determine whether nematode DNA, including RTE1_Sar, may have contaminated the library preparation of the shrew genomic DNA, we assessed the general assembly quality and tested for any contamination from other organisms. If the *S. araneus* genome assembly contains other genes from nematodes with RTE1_Sar, it is more likely that RTE1_Sar is also a contaminant of the genome.

We used BUSCO [18] to determine the assembly quality of *S. araneus* by testing for the presence of conserved genes. The assembly had a completeness score of 88% (Table S1), above the average of vertebrates in Ensembl (Figure S1).

To identify potential contamination of the *S. araneus* assembly, we taxonomically partitioned the sequence reads from *S. araneus* with BlobTools [19]. We aligned the sequence reads to the reference assembly with bowtie2 [20] and compared the reads with DIAMOND [21] against a UniProt Reference Proteomes database. Of the 2.5 billion sequence reads, 74% mapped to the *S. araneus* assembly. BlobTools estimated that 7% of the aligned reads are of nematode origin, and 67% are from Chordata (Figure S2). The nematodes are from the distantly related orders of Trichocephalida and Strongylida, respectively Clade I and Clade V [22]. The majority of the reads that mapped to nematode aligned best to species of the genera *Trichuris* or *Trichinella*, both members of the order Trichocephalida (Figure 1). The remainder aligned best to species of the order Strongylida. In contrast to the Trichocephalida reads, those of the Strongylida exclusively aligned to the RTE1_Sar sequence. This finding is consistent with contamination from *Trichuris/Trichinella* genomic DNA but not from the Strongylida species.

**Figure 1:**
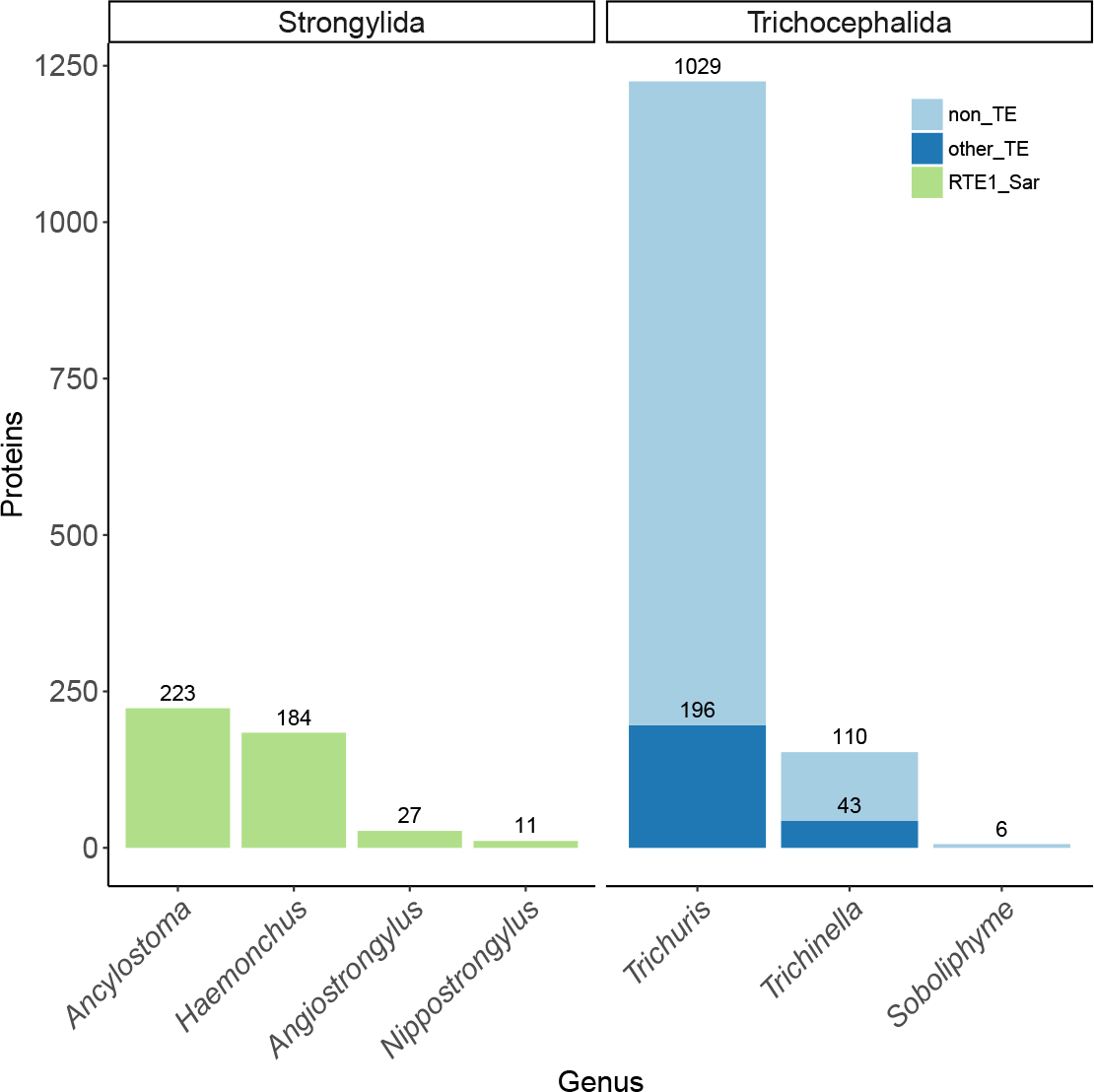
Contamination of the *S. araneus* genome. Proteins are annotated as RTE1_Sar, other TEs and non-TEs. Proteins from Trichocephalida are mostly non-TEs from *Trichuris* and *Trichinella*. Strongylida contribute only RTE1_Sar. Graph only shows genera with more than one protein.

### Test with long reads and read mates reliably identifies horizontal transfer

To further investigate if RTE1_Sar in *S. araneus* results from horizontal transfer, we developed a new, targeted approach that can distinguish horizontal transfer from contamination. Our approach is to reliably determine the origin of reads that contain potential horizontally transferred transposon by matching them to either donor (other) or recipient (self) taxon. If matched to recipient taxon, the transposon is horizontally transferred. If the reads originate from the donor species, they are contaminants. The method includes different approaches depending on the type of data sets are available. We included a positive and negative control. First, we tested if BovB, a transposable element of mammalian origin which we found in textitH. contortus and *O. ochengi*, results from contamination is as such suited as negative control. We suspected that the observation of BovB in these nematode assemblies was a consequence of contamination with host DNA of sheep (*O. aries*) and cattle (*B. taurus*), respectively. PacBio long reads of *H. contortus* were publicly available. We identified reads with BovB, masked them for repeats with RepeatMasker, and mapped them to the two databases as explained above.

95% of BovB-containing long reads (7649 of 8079) originated from a mammal species, not from a nematode. Therefore, the most likely explanation is that BovB transposons in the *H. contortus* assembly are contamination from the host. The absence of long reads for *O. ochengi* prevented us from carrying out a similar test.

We developed three different approaches to determine the origin of reads encoding RTEs, depending on whether the type of sequence data are long reads or paired-end reads, and a reference genome is available. The three approaches were: 1) Long reads: identifying the taxon of origin of the long reads containing one of the RTEs as described above, 2) Reference-based, paired-end reads: aligning the read pairs to the genome assembly, identifying read pairs of which one mate aligned to one of the RTEs and identifying the taxon of origin from the mate, and 3) Non-referencebased, paired-end reads: identifying reads encoding for one of the RTEs and determining the taxon of origin for the non RTE encoding mate (Figure 2). We used BovB as negative control and AviRTE as positive control. PacBio long reads were available for *H. contortus* and Anna’s hummingbird *Calypte anna*, and Illumina read pairs were available for *S. araneus*, *H. contortus* and *T. guttatus*.

**Figure 2:**
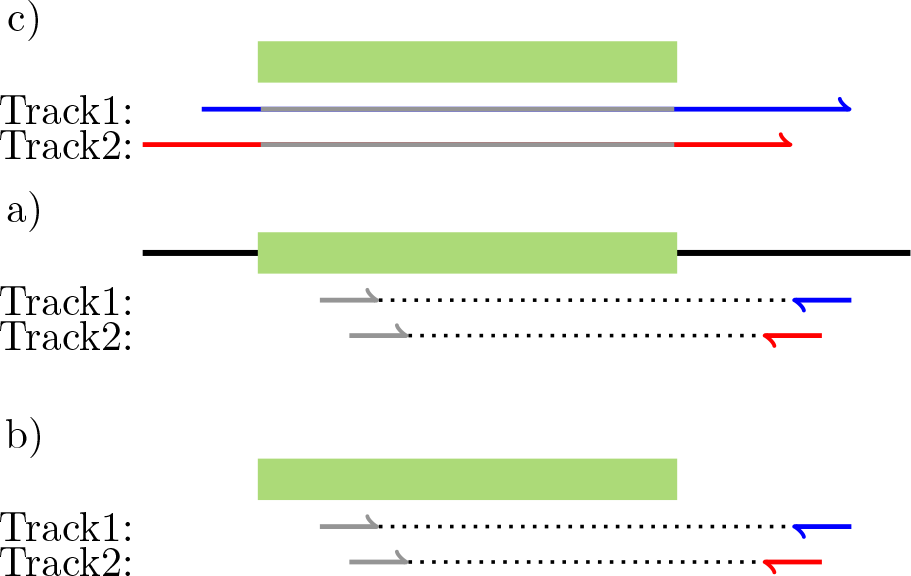
Test for horizontal transfer with read mates and long reads. a) PacBio long reads. b) Reference-based: Illumina read mates aligned to genome assembly. b) Non reference-based: Illumina read mates independently from genome assembly. Track1: horizontal transfer, Track2: contamination. Black line: Genome assembly, green rectangle: RTE, colored arrows: reads (blue arrow: read endogenous, red arrow: read from contaminant.

All approaches agreed that: 1) RTE1_Sar is not contamination in both *S. araneus* and *H. contortus*, 2) our positive control, AviRTE, is not contamination in either bird nor nematode (*Tinamus guttatus/C. anna*; *Loa loa*), and 3) our negative control, BovB in *H. contortus*, is contamination (Figure 3). This shows that these approaches can be used to distinguish between horizontal transfer and contamination. The approaches differ however in the number of informative reads. In all but one case, BovB in *H. contortus*, the reference method reports more reads than the non-reference method. The non reference-based method results in the lowest number of informative reads, which makes it only suited for larger data sets and the reference-based method the preferred approach if long reads are unavailable.

**Figure 3:**
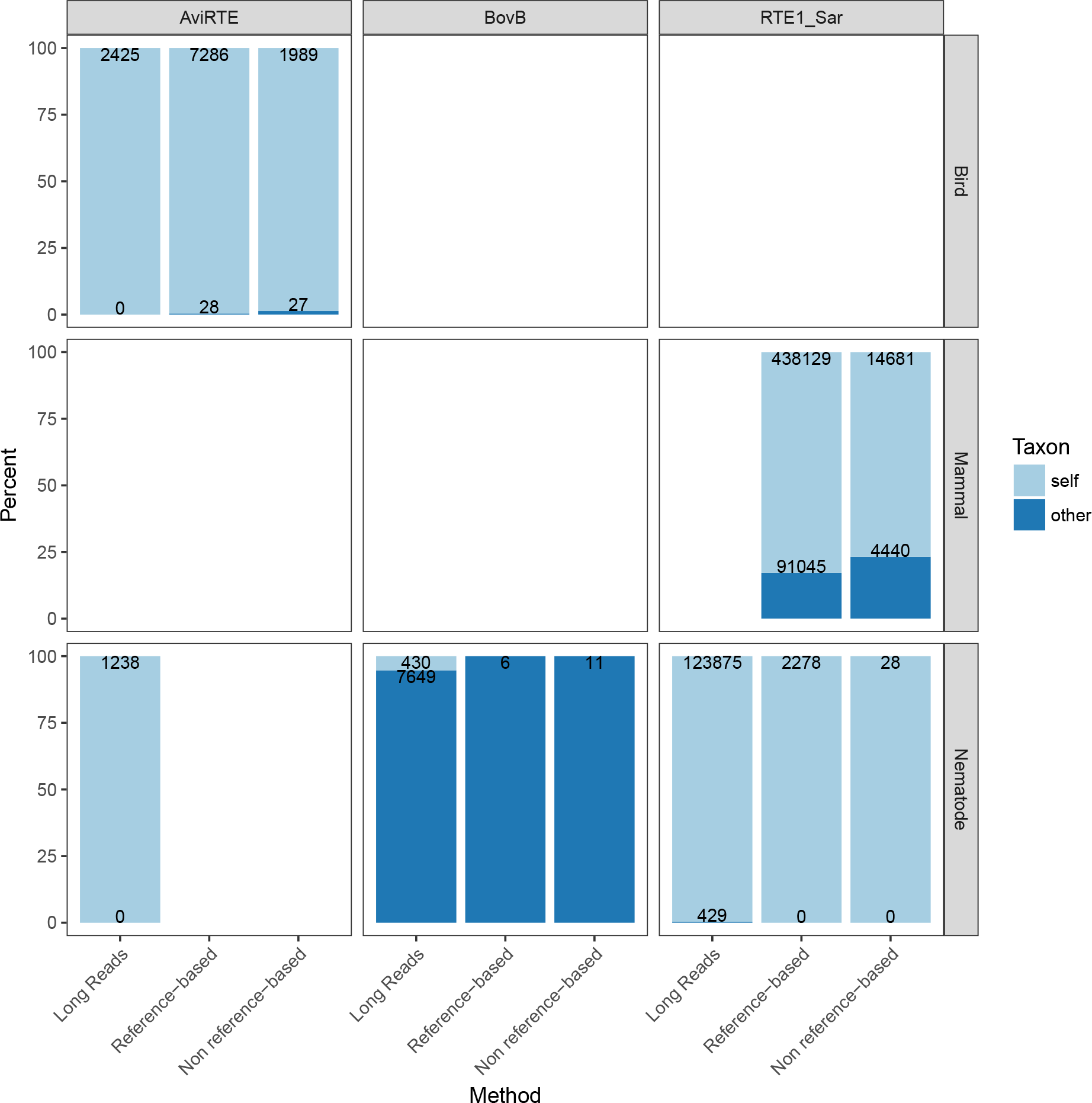
Origin of RTEs in genome assemblies. Y-axis: percent of reads matching to taxon of origin (self) or not (other), x-axis: method used to identify taxon of read mate.

### RTE-1_Sar originates from nematodes

We investigated the origin of RTE1_Sar, a transposon originally found in and named after *S. araneus*. We found RTE1_Sar in nine nematode species, all members of the Trichostrongyloidea superfamily. Within the mammals, our searches showed that RTE1_Sar is found only in *S. araneus*. We have previously shown that mis-assembly of the genome can be responsible for the perceived absence of a gene [23], which can also be the case for a transposon if only few, old or fragmented copies exist. Therefore, we searched for RTE1_Sar in the short sequence reads for species of the order of Eulipotyphla, for which genome projects are available: the European hedgehog *Erinaceus europaeus*, the Hispaniolan solenodon *Solenodon paradoxus woodi* and the star-nosed mole *Condylura cristata*, each representing a different family within the Eulipotyphyla. We found no significant hits (e-value below 1e-10, sequence identity above 75%) in any of these species. This confirms that the taxonomic distribution of RTE1_Sar spans multiple nematode species with divergent host specificities and only one mammal. This strongly suggests that RTE1_Sar arose in nematodes and subsequently transferred to *S. araneus*.

### RTE1_Sar was horizontally transferred from parasitic nematodes to the common shrew

To support the hypothesis of horizontal transfer of RTE1_Sar, we estimated a phylogeny and relative RTE1_Sar copy ages. *S. araneus* RTE1_Sar copies are highly similar to nematode copies (Figure S3), indicating that RTE1_Sar could have been horizontally transferred from nematodes. We estimated the phylogeny of RTE1_Sar. We built consensus sequences from the RTE1_Sar sequences previously identified with BLAST, and built a phylogeny with MrBayes usingRTE1 from *C. elegans* as outgroup. In the resulting phylogeny, RTE1_Sar of *S. araneus* is the sister clade to RTE1_Sar in *H. polygyrus* and *N. brasiliensis*, which are parasites of rodents (Figure 4). To assess the RTE1_Sar repeat landscape and the age of RTE1_Sar copies, we compared the sequences of RTE1_Sar copies to the reference consensus sequences for each species and calculated the Kimura distance with RepeatMasker. The more similar a copy is to the consensus sequence, the shorter is its distance and the younger its age. The Kimura distances of RTE1_Sar copies are shorter in *S. araneus* than in nematodes (Figure 5). Older copies in nematodes have a Kimura distance of up to 50, while the oldest copies in *S. araneus* have a Kimura distance of under 40. We did not observe a sudden replication burst for older copies in *S. araneus*. Most species, including *S. araneus*, have a higher amount of younger copies, especially *A. ceylanicum*, *An. cantonensis*, *An. costaricensis*, and *H. polygyrus*.

**Figure 4:**
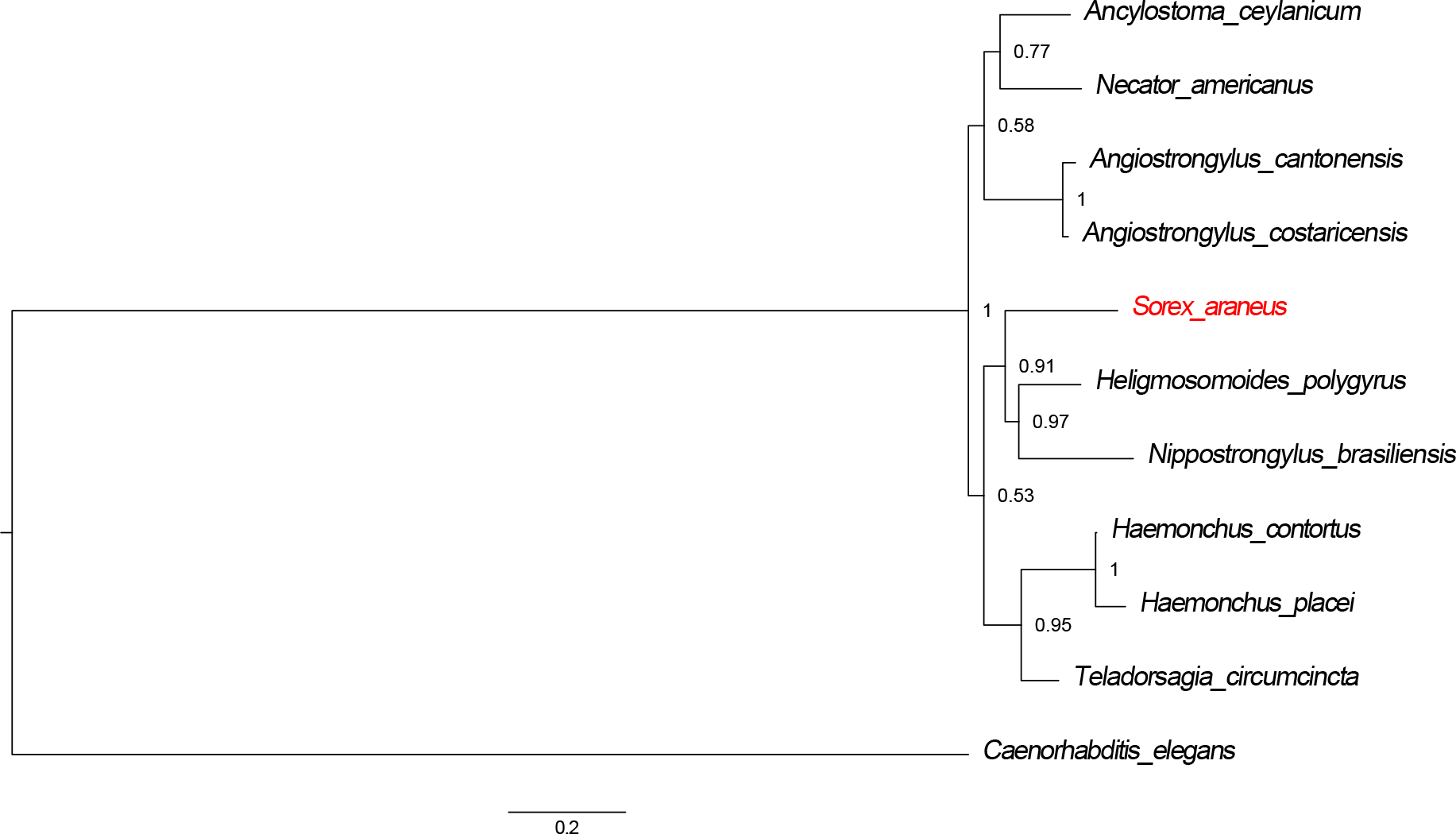
Phylogeny of RTE1_Sar in nematodes and *S. araneus*. RTE1_Sar of *S. araneus* is most closely related to RTE1_Sar in *H. polygyrus* and *N. brasiliensis*, indicating horizontal transfer from an ancestor a closely related species to *S. araneus*. *C. elegans* RTE1 was used as outgroup. Values at nodes represent probability from MrBayes analysis.

**Figure 5:**
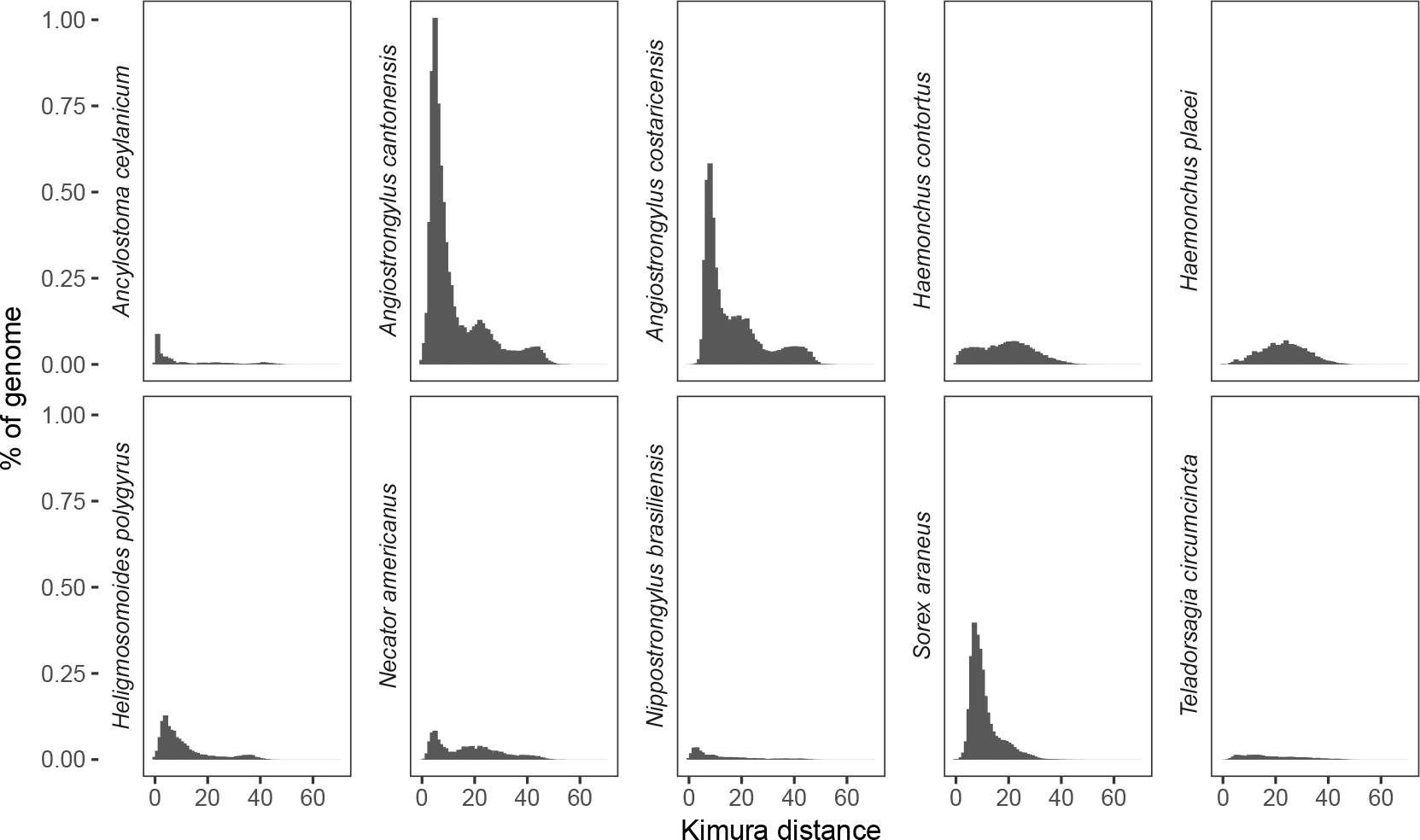
Relative ages of RTE1_Sar copies in their host genomes. Copies in *S. araneus* are younger and much more abundant (total number) than in nematodes. *A. ceylanicum*, *An. cantonensis*, *An. costaricensis* and *N. americanus* present multiple peaks and their RTE1_Sar might have undergone multiple replication bursts.

## Discussion

HTTs occur frequently in metazoa, with 2772 reported cases on HTT-DB [24] (last accessed 5th September 2018). Some species are more prone for HTT than others, such as parasites that might facilitate HTTs [5, 7]. Two cases of horizontal retrotransposon transfer include parasites either directly (AviRTE) or as vector (BovB). The purpose of this study was to identify potential horizontal RTE transfer between parasitic nematodes and their hosts in order to better understand the frequency of HTT between hosts and parasites. We identified a third RTE, RTE1_Sar, that has been horizontally transferred between parasites and their hosts.

RTE1_Sar, although annotated as transposon from *S. araneus*, originates from nematodes. We found that *S. araneus* is the only mammal with RTE1_Sar, and a previous study showed that RTE1_Sar is closer related to insect and nematode RTEs than to mammalian RTEs [4]. This shows that the transposon annotation in Repbase does not necessarily represent the species or lineage of origin, but the species in which the transposon was first identified. This needs to be considered in future studies, especially those investigating horizontal transfers.

Distinguishing between horizontal transfer and contamination is not easy. Careful analyses in the tardigrade [12] and the kleptoplastic sea slugs [25, 26, 27] over-turned earlier pronouncements of horizontal gene transfer. We identified RTE1_Sar in parasitic nematodes, and have shown with subsequent analysis that it is not a contaminant in either nematodes or shrew. We designed a protocol to differentiate between contamination and horizontal transfer with long and short reads. Long reads are more reliable than short reads, because they are continuous and contain more base pairs, which provides more information than short read mates with the same fragment length. In the absence of long reads for *S. araneus*, we used paired-end short reads and validated them with PacBio long read libraries from *H. contortus*, *L. loa* and *C. anna*, including positive and negative controls. The read mate approach provides the same answers as the long read approach, which confirms it as a reliable method to test for contamination when no long reads are available.

*S. araneus* RTE1_Sar is the sister taxon of RTE1_Sar from *H. polygyrus* and *N. brasiliensis*. There are two possibilities for the horizontal transfer of RTE1_Sar into *S. araneus*, either from an unsampled parasitic nematode which is closely related to the two nematode species, or their common ancestor. The transfer happened after the split of the Soricinae and *Erinaceus* lineages which occurred ca. 64 million years ago (mya) [13], since we did not detect the element in *E. europaeus*, the closest related species to *S. araneus* with a sequenced genome.

We have analyzed the relative ages of RTE1_Sar copies across all species to describe the RTE1_Sar landscape and to discover potential replication bursts that might follow horizontal transfers. We did not however observe sudden replication bursts of older copies in the shrew. Replication bursts indicate a horizontal transfer: genomes newly exposed to a transposon would not have any specific defense mechanisms, which allows for faster replication. The absence of replication bursts could be explained by pre-existing defense mechanisms from similar transposons.

We do not know the mechanisms of horizontal transposon transfer, but we suspect that there is a higher chance of HTT between physically or spatially connected species, such as parasites and their hosts. The most encompassing HTT study so far was conducted in insects, and found that transposons are more likely to be transferred to species sharing closer habitat space [28]. We are still lacking truly comprehensive studies encompassing possible transmission routes between classes or even kingdoms. Venner et al. suggested the use of networks to find transposon transmission routes between interacting species, including parasites and pathogens [29]. By knowing the organisms involved in HTT, we will be able to narrow down possible transfer mechanisms. Both viruses and endosymbiontic bacteria have been suggested as vectors [30, 5, 4, 31]. With respect to nematodes, horizontal gene transfers from prokaryotes have been identified [32, 33, 34]. Beneficial plant cell wall-degrading enzymes were transferred from prokaryotes to plant parasitic nematodes [32]. HTT could have equally important impact on a species.

## Conclusion

By understanding horizontal transposon transfers we get one step closer to understanding host-parasite interactions and their consequences on genome evolution. Here, we demonstrated that RTE1_Sar has transferred from parasitic nematodes to *S. araneus*, and we presented a new method to distinguish contamination from horizontal transfer. We confirm, in addition to studies of BovB and AviRTE, that RTEs can jump between species in close associations such as parasites and their hosts. More studies are needed to estimate the frequency of horizontal transfers, and to investigate the diverse effects this can have on the new host, its genome and its interactions with the environment.

## Methods

### RTE detection in genome assemblies of nematodes and mammals

To screen for RTEs of nematodes and mammals in the respective other taxon in order to identify potential horizontal transfers, we used reciprocal similarity searches. We downloaded 10 nematode and 33 mammal RTEs from Repbase Update (version 21.04.14) [14]. We obtained all representative mammalian genomes from RefSeq [16] (98 total), and all nematode genomes from Wormbase Parasite 6 [15] (81 total). We compared the RTE sequences to the genomes with BLASTn [35] (v2.4.0+, blastn -evalue 1e-10) searches, and filtered the results using a length cutoff of 100bp. We extracted the genomic locations of each hit from the genome using Bedtools v2.24.0 [36] while filtering duplicates, and used BLAST to extract the nucleotide sequences. We performed a best hit reciprocal similarity search to compare the extracted sequences back to the Repbase database.

### *S. araneus* genome quality and contamination

To test for genome completeness, we quantified the number of conserved genes present in the assembly of *S. araneus* (SorAra2.0, GCA 000181275.2) using BUSCO v3.0.2 [18].

We used BlobTools v1.0 [21] to detect contamination in the *S. araneus* genome. We used Bowtie2 v2.3.3.1 [20] to align sequence reads to the genome (default settings). The sequence reads were downloaded from the NCBI SRA (Bioproject PRJNA13689). To determine the taxonomy of contamination, we used DIAMOND v0.9.10 [21] and compared the genome assembly against a UniProt Reference Proteomes database [37] (downloaded November 2017, `diamond blastx –max-target-seqs 1 –evalue 1e-25 –sensitive`).

### RTE1_Sar detection in sequence reads of Eulipotyphla

We searched for RTE1_Sar in the sequence reads of three additional Eulipotyphla species. We downloaded the sequence reads of *C. cristata*, *E. europaeus*, and *S. paradoxus woodi* from the NCBI sequence read archive [38] (BioProjects PRJNA74585, PRJNA368679 and PRJNA72447), and the amino acid sequence of the open reading frame (ORF) of RTE1_Sar (RTE1_Sar 1p) from Repbase. We used DIAMOND [21] to identify the ORF in the sequence reads (`diamond blastx –sensitive`). The results were filtered for sequence identity above 75% and e-values below 1e-10.

### Phylogeny

We estimated the phylogeny of the RTE1_Sar sequences across species to understand the relationship of RTE1_Sar in *S. araneus* and nematodes. For each species, we aligned all identified RTE1_Sar sequences with MAFFT v7.310 using the accuracy-oriented method ‘E-INS-i’, and built consensus sequences from the alignments with hmmbuild and hmmemit (HMMER v3.1b2, [39]). We aligned the species’ consensus sequences of RTE1_Sar with MAFFT and built a phylogenetic tree with MrBayes v3.2.6 [40] (GTR model with gamma distribution, 12 * 10^5^ generations run length).

### Relative age distribution

We calculated the Kimura 2-parameter distance (excluding CpG sites) using the package calcDivergenceFromAlign.pl from RepeatMasker for all RTE1_Sar and RTE1 copies to their species specific consensus sequence as estimation of the relative age of each copy compared to the species consensus sequence. We plotted the age distributions with R [41].

### Determination of the taxon of origin of RTE encoding sequence reads

To determine whether sequence reads with RTEs originated from the sequenced organism and not from contamination, we compared them to two respective databases. We used three different protocols, one for long reads and two for paired-end reads.

#### Long reads

We downloaded PacBio long reads for *H. contortus*, *C. anna* and *L. loa* (Bioprojects PRJEB2252, PRJNA289277: 100 pM loading concentration, and PRJNA246086). We searched for the respective RTEs (RTE1_Sar, AviRTE CAn (*C. anna*), BovB from *Bos taurus*) with diamond blastx in the sequence reads (`diamond blastx –more-sensitive`), using the ORFs (open reading frames) of the RTEs. The ORF sequences were aquired from RepBase. The fasta sequences of reads with hits were extracted from the fastq files with seqtk v1.2-r94 (https://github.com/lh3/seqtk) and masked for transposons with RepeatMasker against RepBase (2018). The masked sequences were then blasted (blastn -evalue 1e-10) against two databases to find the origin of the read pair. The databases consisted of the genomes from nematodes and mammals or birds (Refseq, n=97) respectively. The hits were ranked by bitscore, and hits to the genome(s) of the tested species were discarded.

#### Reference-based: Illumina mates aligned to reference assembly

We downloaded paired-end sequencing reads from NCBI for *H. contortus*, *T. guttatus* and *S. araneus* (Bioprojects PRJEB4207, PRJNA212876 and PRJNA13689). We annotated the reference assembly for locations of the respective RTE with RepeatMasker, and produced a bedfile by filtering for alignments with sequence divergence under 20% and longer than 200bp. We aligned the paired-end reads to the genome with Bowtie2 and filtered for reads overlapping the annotated RTE regions with Bedtools intersect. We used Samtools [42] to discard read pairs if both mates overlapped RTE regions, and kept the singletons. The singletons and their mates were screened for transposons in Repbase: singletons reciprocally mapping to the RTE were kept, and their mates masked for all repetitive elements. The mates were then mapped against two databases as described above.

#### Non reference-based: Illumina mates without reference assembly

We downloaded paired-end sequencing reads from NCBI for *H. contortus*, *Tinamus guttatus* and *S. araneus* (Bioprojects PRJEB4207, PRJNA212876 and PRJNA13689) and filtered the sequences for quality with seqtk (seqtk seq -q20). The filtered reads were scanned for the respective RTE (RTE1_Sar in *S. araneus*, AviRTE in *T. guttatus*, and 17 BovBs in *H. contortus* (List of BovBs see Supplemental File2). Mates of reads with reciprocal hits that do not contain the respective RTE were searched (blastn -evalue 1e-03) against the two databases mentioned above.

## Acknowledgement

We are grateful to Dr. Alexander Suh for helpful discussions and suggestions. We thank Dr. Sam Yeaman and Ivan Krukov for their feedback on the manuscript.

## Supplemental Files

### Supplemental File 1

List of genomes.

### Supplemental File 2

List of RTEs.

### Supplemental File 3

**Table S1 and Figures S1-3**. Table S1 contains the BUSCO scores for the *S. araneus* genome assembly.

**Figure S1** shows BUSCO scores for Ensemble vertebrates, **Figure S2** shows the results of the BlobTools analysis for the *S. araneus* genome assembly, and **Figure S3** shows a matrix with sequence identities of RTE1_Sar consensus sequences.

## References

[1] D. Kordis and F. Gubensek. Unusual horizontal transfer of a long interspersed nuclear element between distant vertebrate classes. Proc. Natl. Acad. Sci. U.S.A., 95(18):10704–10709, Sep 1998.

[2] A. M. Walsh, R. D. Kortschak, M. G. Gardner, T. Bertozzi, and D. L. Adelson. Widespread horizontal transfer of retrotransposons. Proc. Natl. Acad. Sci. U.S.A., 110(3):1012–1016, Jan 2013.

[3] A. M. Ivancevic, R. D. Kortschak, T. Bertozzi, and D. L. Adelson. Horizontal transfer of BovB and L1 retrotransposons in eukaryotes. Genome Biol., 19(1):85, 07 2018.

[4] A. Suh, C. C. Witt, J. Menger, K. R. Sadanandan, L. Podsiadlowski, M. Gerth, A. Weigert, J. A. McGuire, J. Mudge, S. V. Edwards, and F. E. Rheindt. Ancient horizontal transfers of retrotransposons between birds and ancestors of human pathogenic nematodes. Nat Commun, 7:11396, Apr 2016.

[5] S. Schaack, C. Gilbert, and C. Feschotte. Promiscuous DNA: horizontal transfer of transposable elements and why it matters for eukaryotic evolution. Trends Ecol. Evol. (Amst.), 25(9):537–546, Sep 2010.

[6] C. Gilbert, S. Schaack, J. K. Pace, P. J. Brindley, and C. Feschotte. A role for host-parasite interactions in the horizontal transfer of transposons across phyla. Nature, 464(7293):1347–1350, Apr 2010.

[7] C. Gilbert and C. Feschotte. Horizontal acquisition of transposable elements and viral sequences: patterns and consequences. Curr. Opin. Genet. Dev., 49:15–24, Apr 2018.

[8] T. Laha, A. Loukas, S. Wattanasatitarpa, J. Somprakhon, N. Kewgrai, P. Sithithaworn, S. Kaewkes, M. Mitreva, and P. J. Brindley. The bandit, a new DNA transposon from a hookworm-possible horizontal genetic transfer between host and parasite. PLoS Negl Trop Dis, 1(1):e35, Sep 2007.

[9] S. Ayarpadikannan and H. S. Kim. The impact of transposable elements in genome evolution and genetic instability and their implications in various diseases. Genomics Inform, 12(3):98–104, Sep 2014.

[10] B. Chenais, A. Caruso, S. Hiard, and N. Casse. The impact of transposable elements on eukaryotic genomes: from genome size increase to genetic adaptation to stressful environments. Gene, 509(1):7– 15, Nov 2012.

[11] J. K. Pace, C. Gilbert, M. S. Clark, and C. Feschotte. Repeated horizontal transfer of a DNA transposon in mammals and other tetrapods. Proc. Natl. Acad. Sci. U.S.A., 105(44):17023–17028, Nov 2008.

[12] G. Koutsovoulos, S. Kumar, D. R. Laetsch, L. Stevens, J. Daub, C. Conlon, H. Maroon, F. Thomas, A. A. Aboobaker, and M. Blaxter. No evidence for extensive horizontal gene transfer in the genome of the tardigrade Hypsibius dujardini. Proc. Natl. Acad. Sci. U.S.A., 113(18):5053–5058, May 2016.

[13] C. J. Douady and E. J. Douzery. Molecular estimation of eulipotyphlan divergence times and the evolution of “Insectivora”. Mol. Phylogenet. Evol., 28(2):285–296, Aug 2003.

[14] W. Bao, K. K. Kojima, and O. Kohany. Repbase Update, a database of repetitive elements in eukaryotic genomes. Mob DNA, 6:11, 2015.

[15] K. L. Howe, B. J. Bolt, M. Shafie, P. Kersey, and M. Berriman. WormBase ParaSite - a comprehensive resource for helminth genomics. Mol. Biochem. Parasitol., 215:2–10, 07 2017.

[16] N. A. O’Leary, M. W. Wright, J. R. Brister, S. Ciufo, D. Haddad, R. McVeigh, B. Rajput, B. Robbertse, B. Smith-White, D. Ako-Adjei, A. Astashyn, A. Badretdin, Y. Bao, O. Blinkova, V. Brover, V. Chetvernin, J. Choi, E. Cox, O. Ermolaeva, C. M. Farrell, T. Goldfarb, T. Gupta, D. Haft, E. Hatcher, W. Hlavina, V. S. Joardar, V. K. Kodali, W. Li, D. Maglott, P. Masterson, K. M. McGarvey, M. R. Murphy, K. O’Neill, S. Pujar, S. H. Rangwala, D. Rausch, L. D. Riddick, C. Schoch, A. Shkeda, S. S. Storz, H. Sun, F. Thibaud-Nissen, I. Tolstoy, R. E. Tully, A. R. Vatsan, C. Wallin, D. Webb, W. Wu, M. J. Landrum, A. Kimchi, T. Tatusova, M. DiCuccio, P. Kitts, T. D. Murphy, and K. D. Pruitt. Reference sequence (RefSeq) database at NCBI: current status, taxonomic expansion, and functional annotation. Nucleic Acids Res., 44(D1):D733–745, Jan 2016.

[17] K. C. Tung, F. C. Hsiao, K. S. Wang, C. H. Yang, and C. H. Lai. Study of the endoparasitic fauna of commensal rats and shrews caught in traditional wet markets in Taichung City, Taiwan. J Microbiol Immunol Infect, 46(2):85–88, Apr 2013.

[18] R. M. Waterhouse, M. Seppey, F. A. Simao, M. Manni, P. Ioannidis, G. Klioutchnikov, E. V. Kriventseva, and E. M. Zdobnov. BUSCO applications from quality assessments to gene prediction and phylogenomics. Mol. Biol. Evol., Dec 2017.

[19] DR Laetsch and ML Blaxter. Blobtools: Interrogation of genome assemblies [version 1; referees: 2 approved with reservations]. F1000Research, 6(1287), 2017.

[20] B. Langmead and S. L. Salzberg. Fast gapped-read alignment with Bowtie 2. Nat. Methods, 9(4):357–359, Mar 2012.

[21] B. Buchfink, C. Xie, and D. H. Huson. Fast and sensitive protein alignment using DIAMOND. Nat. Methods, 12(1):59–60, Jan 2015.

[22] M. L. Blaxter, P. De Ley, J. R. Garey, L. X. Liu, P. Scheldeman, A. Vierstraete, J. R. Vanfleteren, L. Y. Mackey, M. Dorris, L. M. Frisse, J. T. Vida, and W. K. Thomas. A molecular evolutionary framework for the phylum Nematoda. Nature, 392(6671):71–75, Mar 1998.

[23] A. Gilabert, D. M. Curran, S. C. Harvey, and J. D. Wasmuth. Expanding the view on the evolution of the nematode dauer signalling pathways: refinement through gene gain and pathway co-optionx. BMC Genomics, 17:476, 06 2016.

[24] B. R. Dotto, E. L. Carvalho, A. F. Silva, L. F. Duarte Silva, P. M. Pinto, M. F. Ortiz, and G. L. Wallau. HTT-DB: horizontally transferred transposable elements database. Bioinformatics, 31(17):2915–2917, Sep 2015.

[25] C. Rauch, J. d. Vries, S. Rommel, L. E. Rose, C. Woehle, G. Christa, E. M. Laetz, H. Wagele, A. G. Tielens, J. Nickelsen, T. Schumann, P. Jahns, and S. B. Gould. Why It Is Time to Look Beyond Algal Genes in Photosynthetic Slugs. Genome Biol Evol, 7(9):2602–2607, Aug 2015.

[26] D. Bhattacharya, K. N. Pelletreau, D. C. Price, K. E. Sarver, and M. E. Rumpho. Genome analysis of Elysia chlorotica Egg DNA provides no evidence for horizontal gene transfer into the germ line of this Kleptoplastic Mollusc. Mol. Biol. Evol., 30(8):1843–1852, Aug 2013.

[27] H. Wagele, O. Deusch, K. Handeler, R. Martin, V. Schmitt, G. Christa, B. Pinzger, S. B. Gould, T. Dagan, A. Klussmann-Kolb, and W. Martin. Transcriptomic evidence that longevity of acquired plastids in the photosynthetic slugs Elysia timida and Plakobranchus ocellatus does not entail lateral transfer of algal nuclear genes. Mol. Biol. Evol., 28(1):699–706, Jan 2011.

[28] J. Peccoud, V. Loiseau, R. Cordaux, and C. Gilbert. Massive horizontal transfer of transposable elements in insects. Proc. Natl. Acad. Sci. U.S.A., 114(18):4721–4726, 05 2017.

[29] S. Venner, V. Miele, C. Terzian, C. Biemont, V. Daubin, C. Feschotte, and D. Pontier. Ecological networks to unravel the routes to horizontal transposon transfers. PLoS Biol., 15(2):e2001536, 02 2017.

[30] C. Gilbert and R. Cordaux. Viruses as vectors of horizontal transfer of genetic material in eukaryotes. Curr Opin Virol, 25:16–22, 08 2017.

[31] M. F. Ortiz, G. L. Wallau, D. A. Graichen, and E. L. Loreto. An evaluation of the ecological relationship between Drosophila species and their parasitoid wasps as an opportunity for horizontal transposon transfer. Mol. Genet. Genomics, 290(1):67–78, Feb 2015.

[32] E. G. Danchin, M. N. Rosso, P. Vieira, J. de Almeida-Engler, P. M. Coutinho, B. Henrissat, and P. Abad. Multiple lateral gene transfers and duplications have promoted plant parasitism ability in nematodes. Proc. Natl. Acad. Sci. U.S.A., 107(41):17651–17656, Oct 2010.

[33] S. N. McNulty, J. M. Foster, M. Mitreva, J. C. Dunning Hotopp, J. Martin, K. Fischer, B. Wu, P. J. Davis, S. Kumar, N. W. Brattig, B. E. Slatko, G. J. Weil, and P. U. Fischer. Endosymbiont DNA in endobacteria-free filarial nematodes indicates ancient horizontal genetic transfer. PLoS ONE, 5(6):e11029, Jun 2010.

[34] E. M. Schwarz, Y. Hu, I. Antoshechkin, M. M. Miller, P. W. Sternberg, and R. V. Aroian. The genome and transcriptome of the zoonotic hookworm Ancylostoma ceylanicum identify infection-specific gene families. Nat. Genet., 47(4):416–422, Apr 2015.

[35] S. F. Altschul, W. Gish, W. Miller, E. W. Myers, and D. J. Lipman. Basic local alignment search tool. J. Mol. Biol., 215(3):403–410, Oct 1990.

[36] A. R. Quinlan and I. M. Hall. BEDTools: a flexible suite of utilities for comparing genomic features. Bioinformatics, 26(6):841–842, Mar 2010.

[37] T.UniProt Consortium. UniProt: the universal protein knowledgebase. Nucleic Acids Res., 46(5):2699, Mar 2018.

[38] R. Leinonen, H. Sugawara, and M. Shumway. The sequence read archive. Nucleic Acids Res., 39(Database issue):19–21, Jan 2011.

[39] S. R. Eddy. Accelerated Profile HMM Searches. PLoS Comput. Biol., 7(10):e1002195, Oct 2011.

[40] F. Ronquist, M. Teslenko, P. van der Mark, D. L. Ayres, A. Darling, S. Hohna, B. Larget, L. Liu, M. A. Suchard, and J. P. Huelsenbeck. MrBayes 3.2: efficient Bayesian phylogenetic inference and model choice across a large model space. Syst. Biol., 61(3):539–542, May 2012.

[41] R Core Team.R: A Language and Environment for Statistical Computing. R Foundation for Statistical Computing, Vienna, Austria, 2013.

[42] H. Li, B. Handsaker, A. Wysoker, T. Fennell, J. Ruan, N. Homer, G. Marth, G. Abecasis, and R. Durbin. The Sequence Alignment/Map format and SAMtools. Bioinformatics, 25(16):2078– 2079, Aug 2009.

